# Comparing diploid and triploid apples from a diverse collection

**DOI:** 10.1101/2024.07.17.603958

**Authors:** Elaina Greaves, Thomas Davies, Sean Myles, Zoë Migicovsky

## Abstract

Apples (*Malus* X. *domestica* Borkh.) are an economically important fruit species and the focus of continuing breeding efforts around the world. While most apple varieties are diploid, ploidy levels vary across the species, and triploids may be used in breeding despite poor fertility. The impact of ploidy on agricultural traits in apple is not well understood but is an important factor to consider when breeding new apple varieties. Here, we use mean heterozygosity values to categorize 970 apple accessions as diploid or triploid and then contrast apples of varying ploidy levels across 10 agriculturally important traits with sample sizes ranging from 427 to 928 accessions. After correction for multiple testing, we determine that triploids have significantly higher phenolic content. By examining historical release dates for apple varieties, our findings suggest that contemporary breeding programs are primarily releasing diploid varieties, and triploids tend to be older varieties. Ultimately, our results suggest that phenotypic differences between diploids and triploids are subtle and often insignificant indicating that triploids may not provide substantial benefit above diploids to apple breeding programs.

## 1. Introduction

Variation in the number of complete sets of chromosomes in the genome — known as ploidy level — is a widespread phenomenon in plants, and plays an important role in evolution, reproduction, and crop improvement. In plants, varying ploidy has been described, from haploid (1x) cotton^[1]^ to octoploid (8x) strawberry^[2]^, and beyond^[3]^. Ploidy can occur in even numbers of sets (diploid, tetraploid, octoploid) and odd numbers of sets (haploid, triploid). From the possible levels of ploidy, triploidy (3x) is known to produce phenotypic advantages in many plant species. For example, triploid *Populus* have faster growth rates, larger leaves, and are more vigorous than their diploid counterparts^[4]^. While the mechanism underlying improved traits in triploids is not well understood, phenotypic improvements are suspected to be a product of increased organelle production, including chloroplasts, in vegetative tissues^[4,5]^. In crop species, some triploid varieties have been selected for unusual or novel phenotypes, such as high-yield cassava^[6]^ and seedless mandarin^[7]^, however triploid plants generally have decreased fertility^[8]^. Although there are examples of triploidy impacting vegetative growth and fertility traits, the degree to which triploidy produces phenotypic differences across a wide array of fruit traits in agricultural plants remains unclear.

The apple (*Malus* X. *domestica* Borkh.) is an economically important fruit crop and the focus of continuing breeding efforts worldwide. Most apple varieties are diploid, but triploids and tetraploids have also been reported^[9–11]^. Triploids are found among significant commercial varieties (e.g. ‘Ribston Pippin’, ‘Mutsu’, and ‘Jonagold’) and constitute a substantial proportion of commercial apple collections^[12]^. Triploid apple varieties are often cited as producing larger fruit, being more disease resistant, and being more vigorous^[9,13]^. Some contemporary breeding programs intentionally incorporate triploid apple varieties with the aim of harnessing such improvements for the production of novel varieties^[14]^. Despite the inclusion of triploids in apple breeding programs, uneven ploidy causes the production of fewer balanced gametes, generally leading to reduced fertility, which can make them challenging to use effectively in breeding^[15]^. While historical pedigrees suggest that triploids played an important role in producing modern apple varieties, this conflicts with recent genetic analyses. Based on historic apple collections, triploid apples used in apple breeding have rarely produced offspring and have been dubbed breeding “dead ends”^[16]^. Therefore, the benefits of using triploids in apple breeding remain unclear.

Although some have documented improved phenotypes in triploid apple varieties^[9]^, few large and comprehensive comparisons of diploid and triploid apple varieties have been conducted. Quantifying variation in fruit quality across apples of differing ploidy levels not only contributes to our understanding of the impact of ploidy variation on fruit quality, but it can also assist apple breeders in making informed decisions about the use of triploids as potential parents. Here, we use ∼100,000 single nucleotide polymorphisms (SNPs) to differentiate between diploids and triploids. We analyze differences across 10 key agricultural phenotypes and examine apple ploidy as a function of release year, ultimately describing subtle differences in fruit quality and a reduction in triploid in contrast to diploid apple varieties released since 1900.

## 2. Methods

Canada’s Apple Biodiversity Collection (ABC) is an orchard consisting of 1,119 different apple accessions located at the Agriculture and Agri-Food Canada (AAFC) Kentville Research Station in Nova Scotia, Canada. The trees in this collection were initially grafted onto M.9 rootstock in 2011 and then planted in May 2013. The orchard was managed to industry standards. A comprehensive description of the collection is available from Watts et al. (2021)^[17]^.

### Genomic data collection and analysis

The collection of genomic data and identification of SNPs for the apple accessions analyzed in this study was previously described by Migicovsky et al. (2022)^[18]^. Briefly, we performed genotyping-by-sequencing (GBS) using ApeKI and PstI-EcoT22I restriction enzymes and identified 278,231 SNPs across 1,175 apple accessions from the ABC. For this study, we first reduced the genomic dataset, using PLINK (v1.07)^[19,20]^, to only include the 970 *M*. *domestica* accessions harvested in either 2016 or 2017^[17]^. Next, we used a threshold of 0.15 minor allele frequency (MAF) to filter SNPs, resulting in 105,599 SNPs remaining. We removed SNPs with excess heterozygosity using a threshold of 90%, which removed an additional 75 SNPs for a total of 105,524 SNPs. We also generated a secondary SNP file by pruning the SNP set for linkage disequilibrium (LD) using PLINK (v1.07)^[19,20]^ using the command -indep-pairwise 10 3 0.5, resulting in 51,823 LD-pruned SNPs.

We calculated heterozygosity-by-individual using the PLINK command --het and the full SNP set of 105,524 markers before plotting the resulting distribution. Next, we used data from the United States Department of Agriculture (USDA) apple germplasm collection, publicly available from the Germplasm Resources Information Network (GRIN)^[21]^ which determined ploidy of USDA apple accessions using flow cytometry and was previously published in Migicovsky et al. (2021)^[11]^. Based on this information, we calculated the mean heterozygosity-by-individual for accessions labeled as either diploid (2x) or triploid (3x) by GRIN. We plotted these “known” accessions based on ploidy (diploid/triploid) as boxplots and compared the two groups using a Mann-Whitney U-test (wilcox.test) in R version 4.1.0^[22]^. We set a threshold for suspected triploid accessions in the ABC, using the same method outlined in Migicovsky et al. (2021)^[11]^ and labeled all ABC accessions as either diploid or triploid based on this threshold. We also calculated how many ploidy labels assigned using our method differed from the ploidy labels reported by the USDA using flow cytometry measurements.

Using TASSEL^[23]^ we performed principal components analysis (PCA) using the LD-pruned set of 51,823 SNPs and plotted principal components (PCs) 1 and 2, labeling accessions by estimated ploidy level. Finally, we calculated a genome-wide average identity by state (IBS) pairwise identities matrix in PLINK (v1.07)^[19,20]^ using all 105,524 SNPs and the --cluster --matrix options. Based on the IBS matrix, we plotted density distributions for IBS values between diploid-diploid pairs, diploid-triploid pairs, and triploid-triploid pairs. We used a Mann-Whitney U test to determine if the IBS values for triploid-triploid pairs differed from triploid-diploid and diploid-diploid comparisons. All data visualizations were performed using the R package ggplot2 (v3.3.5)^[24]^ in R version 4.1.0^[22]^.

### Phenotype data collection and analysis

For this study, we evaluated the potential consequences of ploidy variation on trait variation using previously published trait (phenotype) data from Watts et al. (2021)^[17]^. Although Watts et al. (2021)^[17]^ report 39 phenotypes, we reduced our dataset to ten phenotypes that we deemed to be of most interest to breeders. Of these ten phenotypes, most were measured in both 2016 and 2017, however, a larger number of apple accessions were measured in 2017. Thus, 2017 data were used when available. When a trait was only measured in 2016, data from this year were used instead. The traits selected were: flowering date, harvest date, time to ripen, weight, acidity, soluble solids content (SSC), the ratio of SSC to acidity, total phenolic content, firmness, and softening during 3 months of cold storage.

A complete description of phenotype data collection methods is available from Watts et al. (2021)^[17]^. Briefly, the flowering date was recorded as the day on which more than 80% of the young branches had king blooms, and the harvest date was recorded as the day when the fruit was deemed ripe and picked from the tree. The time to ripen was calculated as the time between the flowering date and the harvest date. All three phenology measurements were recorded in Julian days.

After harvest, fruit quality traits were collected based on either five (2017) or 10 (2016) fruit. Fruit weight (g) was measured by taking the average weight of harvested fruit. Next, a quarter of each apple was juiced and acidity (g/L) was measured by titrating 1 mL of juice mixed with 0.1 M of NaOH using the 865 Dosimat Plus (Metrohm). Phenolic content was measured using the Folin–Ciocalteu assay and reported in μmol/g fresh fruit. Soluble solids content (SSC) (°Brix) for the juice was also measured using the Pocket Refractometer (Atago, PAL-1). The trait SSC/acidity was calculated by dividing SSC by acidity. Lastly, firmness (kg/cm^2^) was measured by removing apple skin using a vegetable peeler and then using a penetrometer (Guss Fruit Texture Analyser, GS-14).

After harvest, either 10 (2017) or 15 (2016) fruits were placed into cold storage at 3°C to assess how the apples changed during cold storage. After three months of storage, the same methods were used to calculate firmness after storage with a minimum of three and a maximum of 10 apples from each accession. Percent change in firmness (softening) was calculated by subtracting the firmness at harvest from the firmness after storage and then dividing it by the firmness at harvest and multiplying by 100. All the data were analyzed using R version 4.1.0^[22]^ with data visualization performed using the R package ggplot2 (v3.3.5)^[24]^. For all 10 traits of interest, we performed a Mann-Whitney U-test to compare trait values between diploids and triploids. The resulting p values were Bonferroni-corrected (multiplied by 10) to adjust for multiple testing.

Watts et al. (2021)^[17]^ also recorded release year for many of the named apple accessions included in the ABC. Release year is the year in which the named variety was released into commercial production or first mentioned in the literature/media. We incorporated these data into our analyses to determine if there was a change in release year between diploids and triploids. First, we performed a Mann-Whitney U-test to compare release year between diploids and triploids. Lastly, we calculated what percentage of diploid and triploid apples were released before or after 1900. The R code for all analyses is publicly available on GitHub (https://github.com/zoemigicovsky/apple_ploidy).

## 3. Results

### Determining apple ploidy level

To compare diploid and triploid apples, we first evaluated our ability to identify apple ploidy level using the metric heterozygosity-by-individual, calculated based on over 100,000 SNPs genotyped using GBS. After filtering for MAF 0.15, plotting heterozygosity across varieties resulted in a bimodal distribution (Figure 1a). We labeled the mean diploid and triploid heterozygosity-by-individual values based on varieties of known ploidy according to previously published data from the USDA^11]^. Plotting the mean values indicated that the lower peak corresponded to diploid varieties (mean heterozygosity = 0.355) and the peak with higher heterozygosity values corresponded to triploid varieties (mean heterozygosity = 0.471). Based on this distribution, we set a threshold for the suspected diploid/triploid divide of 0.425. We also performed a Mann-Whitney U-test which determined that the two groups of varieties of known ploidy differed (diploid n = 139, triploid n = 48) significantly for heterozygosity by individual values (W = 369, *p* < 1 x 10^-15^) (Figure 1b). Of the varieties labeled by the USDA, we labeled 8 diploids as triploids and 5 triploids as diploids using our threshold. Our method resulted in a total of 12 out of 187 (6.4%) varieties having ploidy labels that did not concur with those previously reported by the USDA GRIN database. Based on our method, 830 (86%) of the apple varieties in the ABC are considered diploid, while 140 (14%) are triploid.

**Figure 1.**
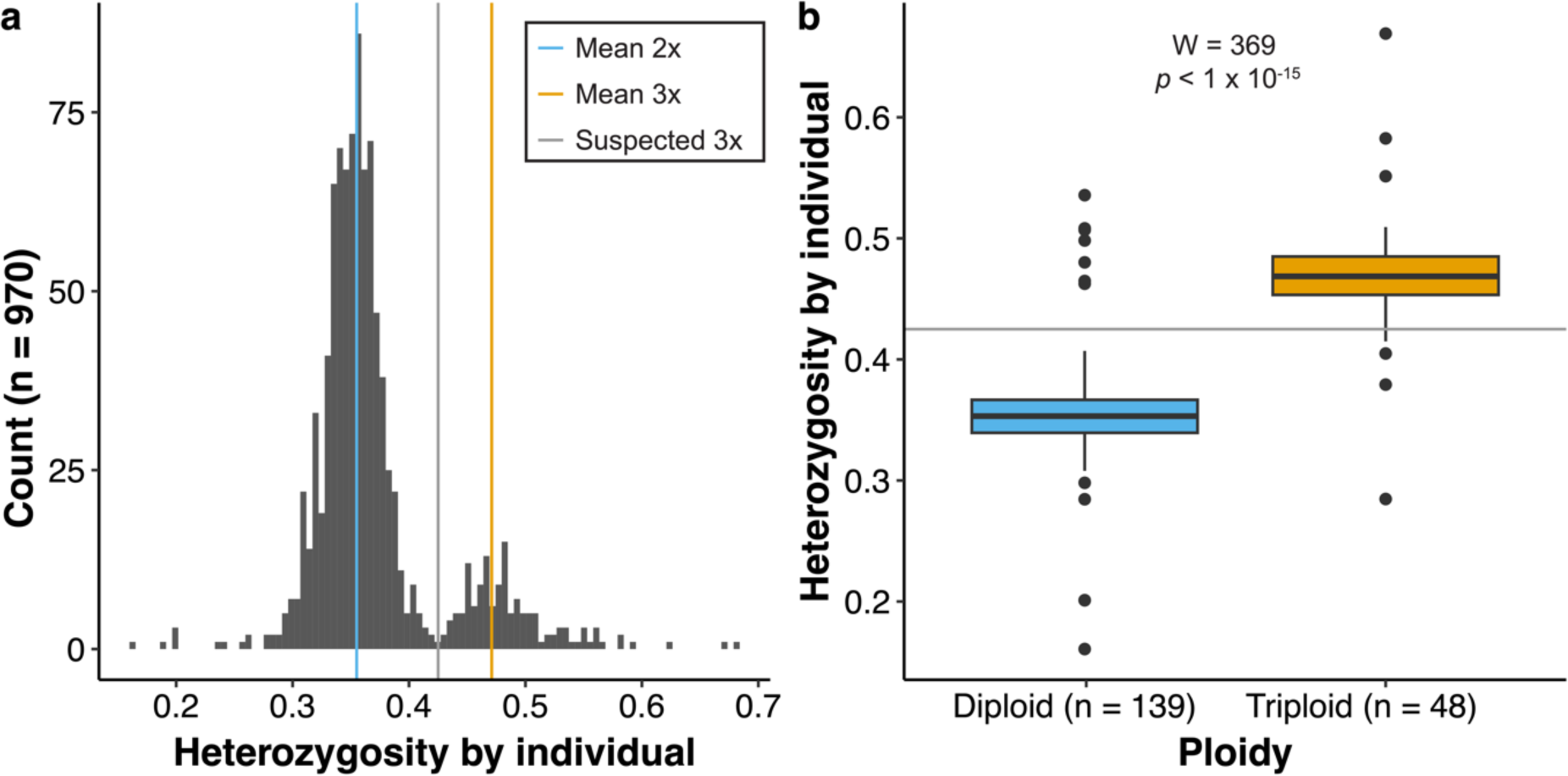
Heterozygosity and ploidy for varieties from Canada’s Apple Biodiversity Collection. (A) Histogram of heterozygosity by individual for all varieties genotyped (n = 970). A blue vertical line indicates the mean value of the diploids (2x), as determined using data from the USDA germplasm collection. The orange vertical line indicates the mean heterozygosity value for triploids (3x) based on data from the USDA. The grey vertical line indicates the threshold used as a cut-off between the two ploidy levels. (B) Boxplots showing heterozygosity by individual for known diploids (blue, n = 139) and triploids (orange, n = 48) with the horizontal grey line indicating the cut-off between the two ploidy levels.

### Relatedness among apple varieties of differing ploidy

After categorizing ABC varieties based on ploidy level (Figure 1), we evaluated relatedness among and across diploid and triploid varieties (Figure 2). First, we performed PCA on 51,823 single nucleotide polymorphisms (SNPs). Principal component (PC) 1 explained 4.51% of the variation in the genomic data, while PC2 explained 3.25% (Figure 2a). Across PC1 and PC2, the values for varieties labeled as both diploids and triploids overlapped. However, the triploid varieties tended to cluster in the middle of the plot with PC1 and PC2 values closer to 0, while the diploid varieties showed a greater range in PC space. For example, no triploid varieties had a PC2 value greater than 40, while some diploid varieties exceeded 60. In addition to PCA, we completed an identity-by-state (IBS) analysis which calculated what proportion of the 105,524 genotyped SNPs shared the same alleles for each pairwise comparison (Figure 2b). We examined the distribution of these IBS values for all pairwise comparisons between two diploid varieties, all pairwise comparisons between two triploid varieties, and all pairwise comparisons between diploid and triploid varieties. Using Mann-Whitney U tests we determined that triploid-triploid comparisons had significantly higher IBS values than those of triploid-diploid (W = 112369979, p < 1 x 10^-15^), or diploid-diploid comparisons (W = 172679637, p-value < 1 x 10^-15^).

**Figure 2.**
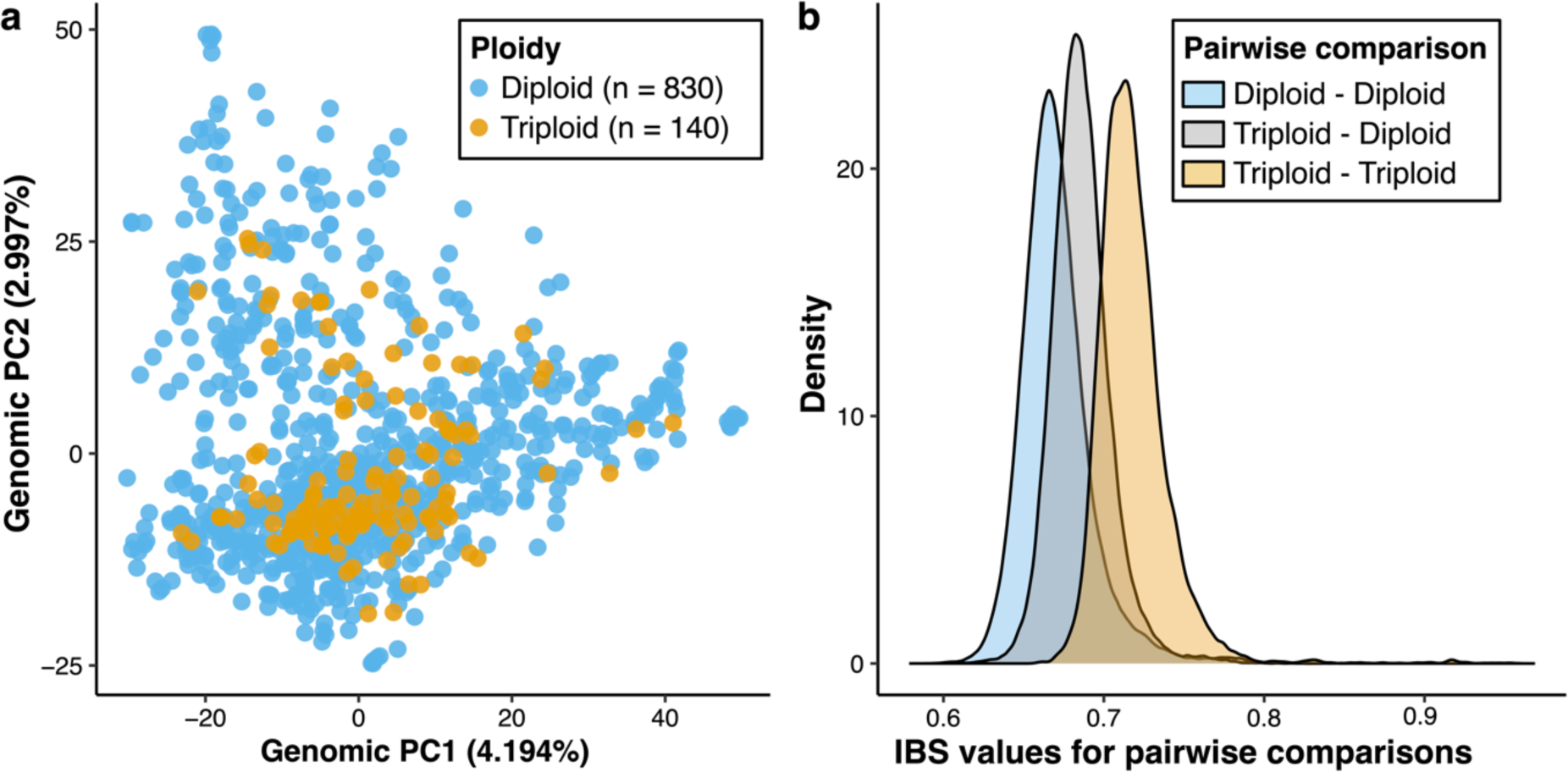
Relatedness among diploid and triploid varieties as determined using genomic data. A) Principal components analysis (PCA) performed using 51,823 single nucleotide polymorphisms (SNPs). Principal component 1 (PC1) and principal component 2 (PC2) are shown with the percentage of variance explained by each PC indicated in parentheses. Each dot indicates a variety and is labeled by the ploidy level, either diploid (blue) or triploid (orange) B) Density plots showing IBS (identity by state) values for each pairwise comparison. IBS values were calculated based on the average proportion of alleles shared at the genotyped SNPs (105,524 SNPs). The blue curve represents all pairwise comparisons between diploids to other diploids, the gray curve represents all comparisons between diploids and triploids, and the orange curve represents all comparisons between triploids to other triploids.

### Trait variation across diploid and triploid apples

In addition to determining how genetically similar diploids and triploids are, we evaluated their phenotypic similarity across ten traits of interest (Figure 3). The total number of apple accessions included for each trait ranged from 427 (phenolic content) to 928 (flowering date). Using a Mann-Whitney U-test to compare diploids and triploids, four traits were significantly different between groups: triploid apples weighed more (W = 27306, p = 0.01075), were firmer (W = 28410, p = 0.04548), had a higher phenolic content (W = 6467, p = 0.002033), and were harvested later (W = 28310, p = 0.03431). However, after Bonferroni correction, the only between group comparison that remained significant was phenolic content (p = 0.02).

**Figure 3.**
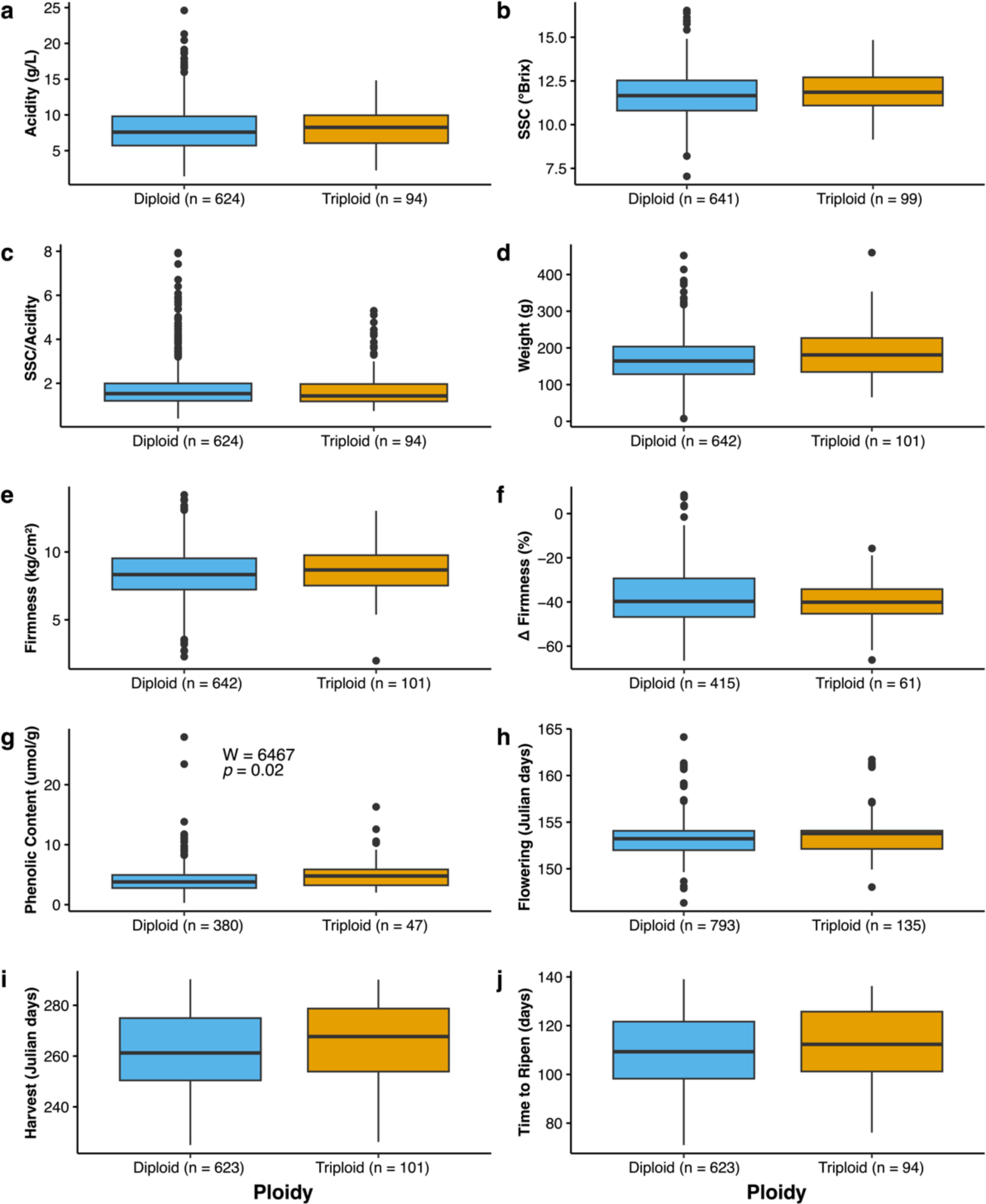
Boxplots showing phenotype values for diploids (blue) and triploids (orange) for the ten traits evaluated. The number of varieties in each group is reported beside the ploidy in all the graphs on the x-axis. A Mann-Whitney U-test was performed to compare trait values between diploids and triploids, and the resulting p-values were Bonferroni-corrected for multiple testing. The test results are shown only for traits where there was a significant difference between ploidy levels after multiple testing corrections. The following traits are shown: a) acidity, b) soluble solid content (SSC), c) SSC divided by acidity, d) weight, e) firmness, f) change in firmness, g) phenolic content, h) flowering date, i) harvest date, and j) time to ripen.

### Comparison of release year for apples of varying ploidy

Lastly, we examined the distribution of release years for diploids and triploids to determine if there was a difference between the groups (Figure 4). The oldest diploid apple in our dataset is ‘Lamb Abbey Pearmain’, which was released in 1804, and diploid apples had an average release year of 1947 and a median release year of 1957. In comparison, the oldest triploid, ‘Ashmead’s Kernel’, was released in 1720 and the average release year for triploids was 1919 while the median was 1947. Overall, release year was significantly later for diploid apples than triploid varieties in our collection (W = 5800, p = 0.048). By comparing varieties from before or after 1900, we determined that 34 out of the 299 diploids with release data were released prior to 1900 (11%), while 10 out of the 32 triploids were released prior to 1900 (31%). Thus, among the apples in our collection with available release date information, a triploid is 2.75 times more likely to be released before 1900 than a diploid apple variety.

**Figure 4.**
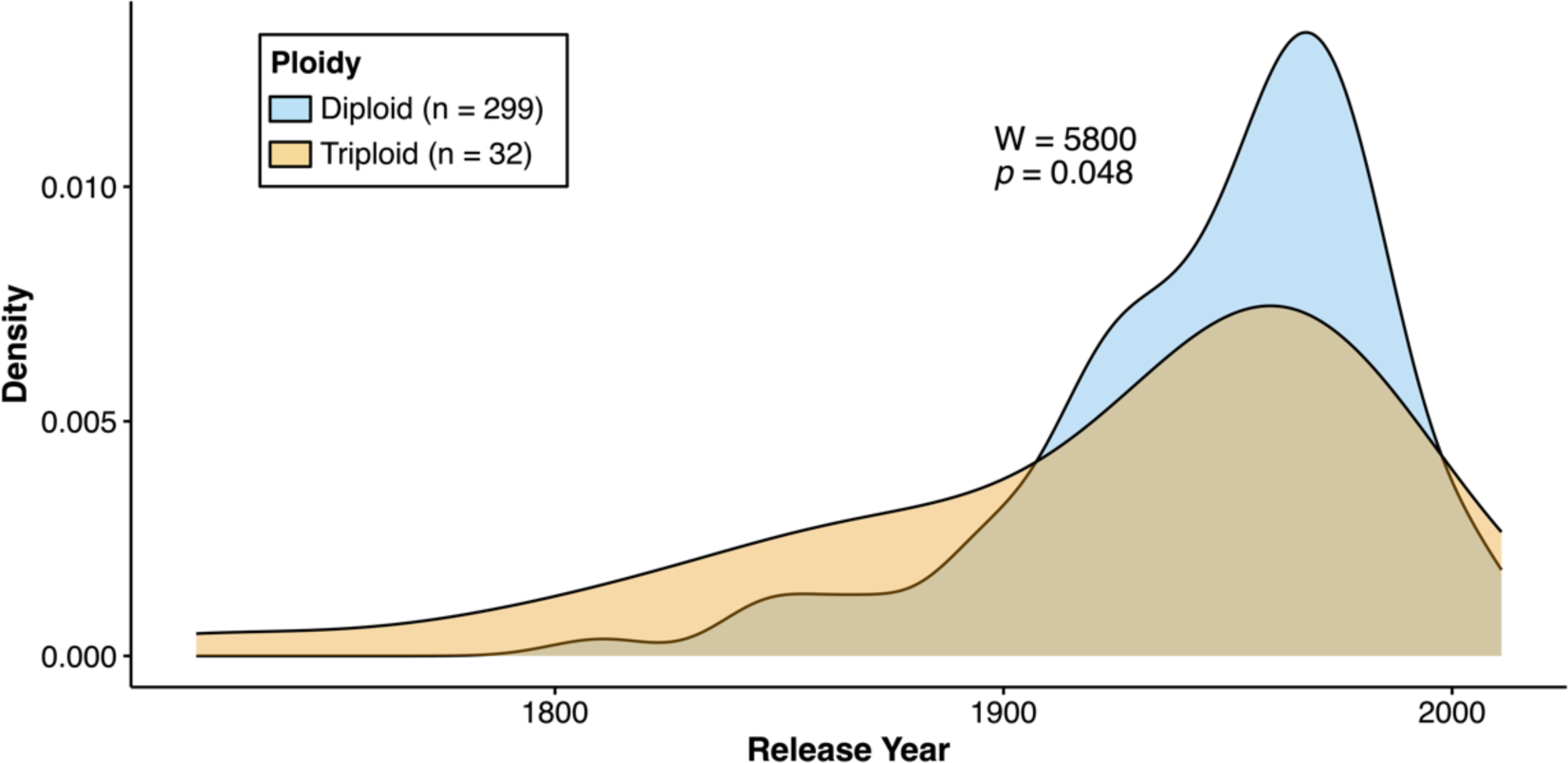
An overlapping density plot showing the date of release (when known) for diploid and triploid apple varieties. The density distribution for diploids (n = 299) is shown in blue and triploids (n = 32) are shown in orange. A Mann-Whitney U-test was performed to compare release year across groups (W = 5800, p = 0.048).

## 4. Discussion

### Genomic variation across diploid and triploid apples

Ploidy level may impact trait variation including vegetative growth and fertility. Apples of varying ploidy have been used in breeding programs and understanding the impact of triploidy on apple fruit quality traits is important for making informed decisions in apple breeding. To make a direct comparison between diploid and triploid apple varieties, we categorized apple accessions from the ABC into diploid and triploid varieties based on heterozygosity (Figure 1a). Heterozygosity is frequently used to differentiate between diploid and triploid in plant samples^[25]^ and has been shown to be a reliable proxy for ploidy in apple^[11,26]^. In our study, less than 6.5% of samples with known ploidy were incorrectly categorized according to USDA flow cytometry information. In these cases, it may be that our method has inaccurately identified ploidy level or that either the variety measured in this study or previous work was mislabeled. However, the high degree of accuracy provides further support for the use of heterozygosity thresholding as a cost-effective and efficient method for categorizing diploids and triploids using GBS^[25,26]^. Indeed, for samples in which flow cytometry ploidy data are available, heterozygosity differs significantly between diploids and triploids (Figure 1b), similar to previous work in kiwifruit (*Actinidia kolomikta*) and aspen (*Populus tremuloides*)^[25,27]^.

Although this analysis supports use of overall sample heterozygosity for accurately distinguishing between diploid and triploid varieties, previous work determined it is unable to distinguish between diploid and tetraploid apple varieties^[11]^, which exist in apple populations^[9,10,14]^ including the USDA apple germplasm collection^[11]^ which largely overlaps with the ABC. Migicovsky et al. (2021)^[11]^ found no significant difference between diploids and tetraploids, indicating that tetraploids in the collection were likely to be primarily autotetraploids. Although some tetraploid apples have been documented to have phenotypes that are significantly different from diploid counterparts^[28,29]^, the impact of tetraploids in the current study is unclear. In addition, the bioinformatics pipeline^[18]^ used to identify SNPs removed SNPs that were not biallelic, therefore removing triallelic SNPs. As a result, triploid diversity was reduced, and informative genetic markers were not included in the present analysis. In summary, although the heterozygosity thresholding used here appears to be an accurate proxy, future work could use flow cytometry to more accurately identify ploidy level prior to comparison of trait variation across apple varieties of varying ploidies.

When examining diploid and triploid apples across the first and second genomic PC, triploids show a smaller range of variability in comparison to diploids (Figure 2a). This may be partially attributable to the smaller size of the triploid samples (N=140) in comparison to the diploids (N=830). It is possible that with a larger group of triploids or a larger amount of genetic information triploids would capture a similar genomic range. However, the ABC is among the largest and most diverse apple collections in the world and our PC analysis suggests that diploid apple varieties capture more genetic variation than triploids but do not significantly differ overall. Future work should consider additional genetic information such as the inclusion of triallelic SNPs or whole genome sequencing to more accurately assess the genetic differences between apples of differing ploidies.

The triploid apple varieties measured in this study form a smaller group that captures less genetic variability within the larger population (Figure 2a) and this finding is supported by our IBS analysis (Figure 2b). IBS values are significantly higher in triploid-triploid comparisons than they are between diploid-diploid or triploid-diploid comparisons. Thus, on average, two triploid varieties will be more closely related to each other than two diploids, or a triploid to a diploid from the collection. This result is primarily a function of comparing the triploids, a relatively small group, to a larger and highly diverse diploid population of apples. Although triploids are significantly more similar to one another than any other comparison in the present study, all three IBS distributions show considerable overlap, indicating that, for example, some diploids are as similar to triploids as some triploids are to each other.

### Trait variation across diploid and triploid apples

In addition to a genomic comparison between diploid and triploid apples, we leveraged existing trait data published for the ABC^[17]^ to compare phenotypic differences between apples of varying ploidies, identifying subtle but insignificant differences in most cases (Figure 3). Across the 10 traits examined, only phenolic content differs significantly between groups after Bonferroni correction for multiple testing (Figure 3g). Prior to correcting for multiple testing, there were measurable differences in fruit weight, fruit firmness, and harvest date between groups, indicating that these differences are subtle and not statistically significant. Previous work has reported that triploid apple varieties differ from diploid apples, for instance having larger fruit^[9]^, improved disease resistance, and greater vigor^[13]^. It has been suggested that these trait improvements have caused selection for triploid apple varieties in the past^[14,30]^. Although there are some accounts of triploid varieties producing larger^[9]^ and firmer fruit^[31]^, after multiple test corrections, we did not detect significant differences between diploid and triploid apples for these traits. However, the trends observed between diploids and triploids in the present work, while statistically insignificant, agree with previous findings. Our findings suggest that across a large and diverse germplasm collection, apples of varying ploidy do not significantly differ across numerous agriculturally important traits. Taken together, this raises questions about the overall utility of selective or preferred use of triploids in apple breeding, particularly given the reduced fertility of triploids^[8,32]^.

Across the ten traits compared between diploids and triploids, the only trait that significantly differed between groups was phenolic content, which was higher in triploid varieties (Figure 3g). To date, there are conflicting reports on the influence of triploidy on the production of fruit metabolites. For example, some comparisons between diploids and triploids from apple breeding programs have noted no differences in the production of fruit metabolites^[9]^. However, a recent investigation of acidity in cider apples found that triploid cider apples were higher in titratable acidity on average than their diploid counterparts^[33]^. Increased acidity in triploids was suspected to be due to the extra copies of genes related to acid production, particularly those at the malic acid *Ma1* locus^[33]^. Similarly, the increase in phenolic content observed here may be a product of increased copies of genes in the biosynthetic pathways that produce phenolic compounds in triploids^[34,35]^. Although this may explain the subtle but significant difference in phenolic content across apples of varying ploidy, it is worth noting that, for example, SSC did not significantly increase in triploid apple fruits despite additional copies of genes responsible for SSC production. Given the measurable increase in phenolic content between the ploidies, triploids may be useful in the breeding of cider apples in the future, particularly if they also have higher acid production^[33]^.

Finally, while the phenotype data from the ABC population are extensive^[17]^, there may be phenotypes such as leaf shape and size, stomatal size, plant height, and plant vigor that differ between diploids and triploids, as seen in other tree species^[4,36,37]^. In the future, a more detailed comparison of plant phenotypes may uncover important phenotypic differences attributable to ploidy.

### Variation in release when comparing diploid and triploid apples

Although we only detect subtle differences in trait variation between diploid and triploid apples, triploid apples have been commercially released and grown for over 200 years (Figure 4). Indeed, the earliest apple release recorded across the 331 varieties examined here was a triploid released 84 years prior to the earliest diploid. That said, these findings are based on previously reported release dates^[17]^ and nearly two thirds of the collection consisted of either non-commercial apples, or those for which release date could not be reliably identified.

The triploid varieties examined here had on average significantly earlier release years than diploid varieties (Figure 4). This finding is supported by recent genetic analyses of historical pedigrees which revealed that triploids have typically been breeding “dead ends”^[16]^. Our work provides further evidence that contemporary breeding programs are primarily releasing diploid apple varieties and that in contrast to diploids, triploid varieties were nearly three times as likely to be released prior to 1900. Given the decreased fertility of triploids^[8,15]^, it is likely that the difference in release date seen here is at least partially attributable to difficulties related to the use of breeding with triploid apples.

## 5. Conclusions

By examining genomic and trait variation across hundreds of unique apple varieties, we were able to compare diploid and triploid apples. We provide further evidence that heterozygosity can be used as a reliable method for estimating ploidy level. Our work suggests that across diverse apple germplasm, triploids are not significantly different from diploids for nine agriculturally important traits. However, we did identify significantly higher phenolic content in triploid varieties, indicating that although differences between diploids and triploids are subtle, there is still measurable variation that may be desirable to harness in certain contexts, such as breeding cider apples. Our work suggests that commercial breeding programs are typically releasing diploid varieties, and the commercial release of triploid apples has decreased since 1900. In summary, these results improve our understanding of the potential impact of triploidy on apple breeding, and call into question the overall utility of including triploids in apple breeding programs.

## 6. Supplementary Material

**Table S1. Assigned apple ploidy for 970 apple accessions examined in this study.** Each accession is linked with the reported ploidy information from the USDA, previously published in Migicovsky et al. 2021)^[11]^, when available. The calculated heterozygosity value and assigned ploidy level are also reported.

## 7. Author contributions

The authors confirm contribution to the paper as follows: study conception and design: Greaves E, Migicovsky Z, Myles S; analysis and interpretation of results: Greaves E, Migicovsky Z, Myles S, Davies T; draft manuscript preparation: Greaves E, Davies T, Migicovsky Z. All authors reviewed the results and approved the final version of the manuscript.

## 8. Acknowledgements

We thank Sophie Watts (Dalhousie University) for their useful input on this article. ZM was supported by funding from the Canada Research Chairs program and the Natural Sciences and Engineering Research Council of Canada (NSERC).

## 10. Data availability

All data and code are publicly available on GitHub (https://github.com/zoemigicovsky/apple_ploidy).

